# Dopamine regulates decision thresholds in human reinforcement learning

**DOI:** 10.1101/2022.09.29.509499

**Authors:** Karima Chakroun, Antonius Wiehler, Ben Wagner, David Mathar, Florian Ganzer, Thilo vanEimeren, Tobias Sommer, Jan Peters

## Abstract

Dopamine fundamentally contributes to reinforcement learning by encoding prediction errors, deviations of an outcome from expectation. Prediction error coding in dopaminergic regions in human functional neuroimaging studies is well replicated. In contrast, replications of behavioral and neural effects of pharmacological modulations of the dopamine system in human reinforcement learning are scarce. Additionally, dopamine contributes to action selection, but direct evidence and process-specific accounts in human reinforcement learning are lacking. Here we examined dopaminergic mechanisms underlying human reinforcement learning in a within-subjects pharmacological approach in male human volunteers (n=31, within-subjects design; Placebo, 150mg L-dopa, 2mg Haloperidol) in combination with functional magnetic resonance imaging and a stationary reinforcement learning task. We had two aims. First, we aimed to replicate previously reported beneficial effects of L-dopa vs. Haloperidol on reinforcement learning from gains. This replication was not successful. We observed no performance benefit of L-Dopa vs. Haloperidol, and no evidence for alterations in neural prediction error signaling. In contrast, Bayesian analyses provided moderate evidence in favor of the null hypothesis. This unsuccessful replication is likely at least partly due to a number of differences in experimental design. Second, using combined reinforcement learning drift diffusion models, we tested the recent proposal that dopamine contributes to action selection by regulating decision thresholds. Model comparison revealed that the data were best accounted for by a reinforcement learning drift diffusion model with separate learning rates for positive and negative prediction errors. The model accounted for both reductions in RTs and increases in accuracy over the course of learning. The only parameter showing robust drug effects was the boundary separation parameter, which revealed reduced decision thresholds under both L-Dopa and Haloperidol, compared to Placebo, and the degree of threshold reduction accounted for individual differences in RTs between conditions. Results are in line with the idea that striatal dopamine regulates decision thresholds during action selection, and that lower dosages of D2 receptor antagonists increase striatal DA release via an inhibition of autoreceptor-mediated feedback mechanisms.

## Introduction

The neurotransmitter dopamine (DA) plays a central role in a range of cognitive processes, including cognitive control^1^, reinforcement learning^2^ and decision-making^3^. Phasic responses of midbrain dopamine neurons encode reward prediction errors, the discrepancy between obtained and expected reward^2^. Prediction error signals play a central role in formal reinforcement learning theory^4,5^. On a neural level, positive prediction errors are thought to be signaled by phasic burst firing of DA neurons^2^, predominantly activating low affinity striatal D1 receptors in the direct pathway that facilitates *go* learning^6–8^. In contrast, negative prediction errors are thought to be signaled by phasic dips of DA neuron firing rates below baseline^2^, predominantly affecting high affinity striatal D2 receptors in the indirect pathway that facilitates *no go* learning^6–8^.

Target regions of midbrain dopaminergic projections^9^ in dorsal and ventral striatum reliably exhibit activation patterns in functional neuroimaging studies that correspond to reward prediction error coding^10,11^. Likewise, animal work has shown a causal role of dopamine neuron signaling in reinforcement learning^12^. On the other hand, causal evidence for a role of DA in human reinforcement learning is primarily based on pharmacological work and studies in patients with dysfunctions in the dopamine system, e.g. Parkinson’s disease (PD).

More generally, contributions of dopamine to reinforcement learning have focused on two aspects, learning (value updating) and performance (action selection)^13,14^. With respect to learning, behavioral and neural effects of pharmacological manipulation of DA appear heterogeneous, replications of specific effects are scarce^15^ and many studies suffer from small sample sizes^15^. Elevation of dopamine transmission via the DA precursor L-Dopa has been suggested to improve in particular *go* learning in healthy subjects^16,17^ and PD patients^7,18^. Such effects might be driven by enhanced neural reward prediction error responses under L-Dopa^15–17^. However, other studies did not find increased striatal reward prediction error responses following L-dopa administration^19,20^, and one study even observed increased punishment-related striatal responses^21^. Other evidence suggests blunted prediction error responses following L-dopa administration in PD patients^22^. Conversely, D2 receptor antagonists have sometimes been reported to impair reinforcement learning^17^, whereas other studies have reported no effects^23^, or effects restricted to post-learning decision-making^24,25^. Interpretation of D2 receptor antagonist effects are complicated by the fact that lower dosages predominantly affect presynaptic D2 autoreceptors^26^, which, via an inhibition of negative feedback^26^, likely increases (rather than decreases) striatal DA release^27–31^. In line with this idea, lower dosages of D2 receptor antagonists in some cases improve learning from positive feedback^8^ (similar to some reported effects of L-Dopa^17^), enhance prediction error signaling^24^ and increase overall stimulus-locked striatal responses^32^. Higher dosages might lead to the reverse effects^33^. This is consistent with attenuated *go* learning and reduced striatal prediction error responses in schizophrenia patients that receive antipsychotics^13^.

But dopamine may also critically contribute to action selection *per se*. One account suggests that increased striatal DA availability during choice increases activation in the striatal *go* pathway, and reduces activation in the *no go* pathway^13,14^, thereby facilitating action initiation vs. inhibition. This resonates with accounts that emphasize a role of DA in regulating response vigor^34–37^. This account is conceptually related to a recent proposal that striatal DA regulates decision thresholds during action selection^38^, which is also supported by basal ganglia circuit models^39^. It is also related to theoretical accounts emphasizing a role for DA in encoding the (subjective) precision of actions and/or policies^40–42^. Decision thresholds play a central role in sequential sampling models^43^ such as the drift diffusion model^44^. In these models, choice behavior as arises from a noisy evidence accumulation process that terminates as soon as the accumulated evidence exceeds a threshold. Decision threshold adjustments are thought to be at least in part regulated by basal ganglia circuits^39,45–51^. In line with a more specific role of DA^38,39^, rodent response time distributions following amphetamine infusion in the striatum change in a manner that is consistent with a threshold reduction^52^. Likewise, increased DA availability in mice increases response rates in the absence of learning^53^, again consistent with a threshold reduction. In humans, administration of the catecholamine precursor tyrosine reduces decision thresholds across different value-based decision-making tasks^54^, and the DA agonist ropinirole reduces decision thresholds during inhibition^55^. In contrast, Bromocriptine (a DA agonist) does not affect thresholds during perceptual decision-making^56^, suggesting that such effects might be task-dependent. Finally, gambling disorder, a putatively hyperdopaminergic disorder^57^, is associated with altered adjustment of decision thresholds over the course of learning^58^. However, direct causal evidence for a role of DA in regulating decision-thresholds in human reinforcement learning is still lacking.

The present study is part of a larger project from which a different learning task has been published previously^19^. The study was initially conceptualized as a replication of the gain condition of Pessiglione et al.^17^. In that study, participants receiving L-dopa (n=13) showed improved learning from rewards, and enhanced striatal coding of positive vs. negative prediction errors, compared to participants receiving Haloperidol (n=13). We scanned a larger sample (n=31) using fMRI, and employed a within-subjects design using slightly higher drug dosages (L-Dopa: 150mg, Haloperidol: 2mg). Because the primary behavioral effect in the original study was observed in the gain condition, the loss and neutral conditions from the original study were omitted here. Note that in particular the isolation of the gain condition likely contributed to our unsuccessful replication of behavioral and neural effects (see discussion).

Recent developments of combined reinforcement learning drift diffusion models^59–63^ (RLDDMs), then allowed us to jointly test DA accounts of learning and action selection within the same data set. First, we aimed to test (and replicate) previously reported effects of DA on reinforcement learning and associated neural prediction error responses^17^. Second, we directly tested a potential role of DA in regulating decision thresholds during reinforcement learning^38,52,54^ by leveraging hierarchical Bayesian RLDDMs, and by directly testing for drug effects on model parameters in a combined model across all three drug conditions.

## Methods

### General procedure

Data were obtained in the context of a larger multi-day combined pharmacological fMRI study^19^ with four testing sessions, performed on separate days. Day 1 consisted of a behavioral testing session, where working memory data and questionnaire data were obtained. On days 2 – 4 (each performed exactly 1 week apart), participants received either Placebo, L-Dopa or Haloperidol (in counterbalanced order, see below for details) and then performed two tasks while brain activity was measured using fMRI. The first task was a restless four-armed bandit task measuring exploration/exploitation behavior. Data from this task have been reported elsewhere^19^. Following a short break, participants then completed the stationary reinforcement learning task based on a previous study^17^, which is reported here. All study procedures were approved by the local ethics committee (Hamburg Board of Physicians) and participants provided informed written constent prior to participation.

### Drug administration

Participants performed three fMRI sessions (in counterbalanced order) under three drug conditions: Placebo, L-dopa (150mg) and Haloperidol (2mg). They arrived in the lab 2.5hrs prior to commencement of fMRI scanning. Upon arrival, they received a first pill containing either 2mg haloperidol or placebo (maize starch). Exactly two hours later, subjects received a second pill containing either Madopar (150mg L-dopa + 37.5 mg benserazide) or Placebo. That is, during the Placebo session, participants received maize starch / maize starch. During the Haloperidol session, they received Haloperidol / maize starch, and during the L-dopa session, they received maize starch / L-dopa.

Half an hour after ingesting the second pill, participants entered the fMRI scanner, where they first performed a restless four-armed bandit task reported elsewhere^19^. Following a short break inside the scanner, they then performed 60 trials of the stationary reinforcement learning task reported here.

Physiological parameters, well-being, potential side effects and mood where assessed throughout each testing session^19^. No participant reported any side effects.

### Reinforcement learning task

During each fMRI session, participants performed a simple stationary reinforcement learning task (see Figure 1) based on a previous study^17^. The task involved two pairs of fractal images (n=30 trials per pair). Per pair, one stimulus was associated with a reinforcement rate of 80% (“optimal” stimulus) whereas the other was associated with a reinforcement rate of 20% (“suboptimal” stimulus). On each trial, options were randomly assigned to the left/right side of the screen, and trials from the two options pairs were presented in randomized order. Following presentation of the two options, participants had a maximum of 3 seconds to log their selection via a button press (see Figure 1). Participants received binary feedback, either in the form the display of a 1€ coin (*reward* feedback, see Figure 1) or as a crossed 1€ coin (*no reward* feedback). Jitters of variable duration (2-6sec, uniformly distributed) were included following presentation of the selection feedback and following presentation of the reward feedback (see Figure 1). Prior to scanning, participants performed a short practice version of the task in order to familiarize themselves with the procedure.

**Figure 1.**
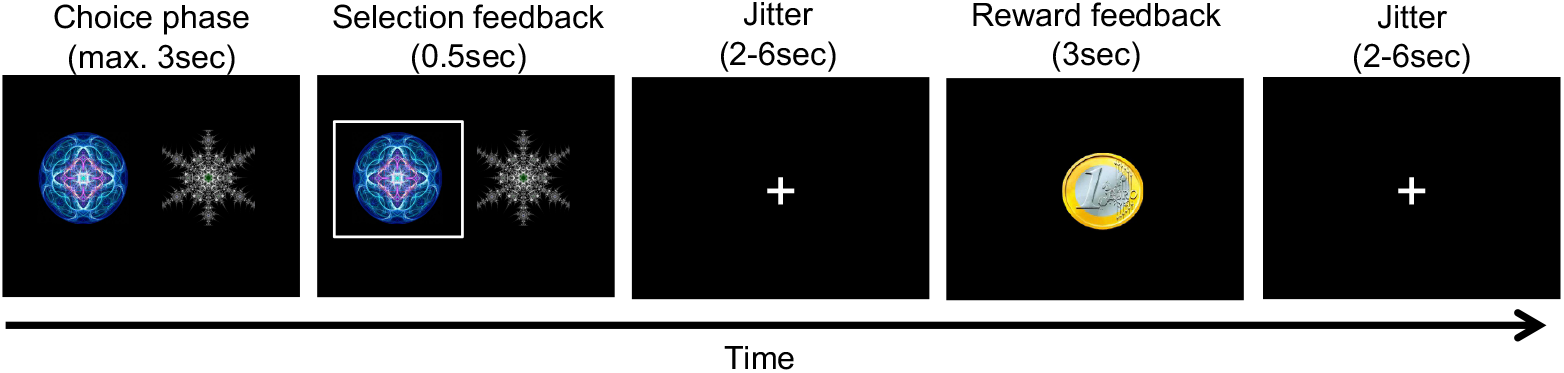
Illustration of a single trial from the reinforcement learning task. Stimuli were presented for a maximum of 3sec, during which participants were free to make their selection. The selection was then highlighted for 500ms, followed by a jitter of variable duration (2-6sec, uniform distribution). Reward feedback was then presented for 3sec, followed by another jitter of variable duration (2-6sec, uniform distribution). Stimuli consisted of two pairs of abstract fractal images (80% vs. 20% reinforcement rate), which were presented in randomized order, and participants completed 30 trials per pair.

### Statistical analyses

Drug effects on model-free performance measures and fMRI parameter estimates extracted at specific peaks were analyzed via Bayesian repeated measures ANOVAs using the JASP software package (Version 0.16.3)^64^.

### Q-learning model

We applied a simple Q-learning model^4^ to formally model the learning process. Here, participants are assumed to update the value (Q-value, Eq. 1) of the chosen option based on the reward prediction error *δ* that is computed on each trial as the difference between the obtained reward and the expected reward. The degree of updating is regulated by learning rate *η* (Eq. 2):

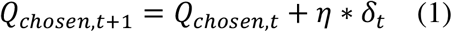

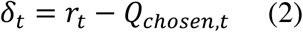

Q-values of unchosen actions remain unchanged. All Q-values were initialized with values of .5. As learning from positive and negative reinforcement is thought to depend on distinct striatal circuits^7,65^, we compared models with both a single learning rate (δ) and dual learning rates (δ_+_, δ_-_) for positive vs. negative prediction errors. Learning rates were estimated in standard normal space [-7, 7] and back-transformed to the interval [0, 1] via the inverse cumulative normal distribution function.

### Reinforcement learning drift diffusion models (RLDDMs)

We used combined reinforcement learning drift diffusion models (RLDDMs)^59,60,66^ to link participants choices and response times (RTs) to the learning model. In the drift diffusion model, choices arise from a noisy evidence accumulation process^43,44^ that terminates as soon as the accumulated evidence exceeds a threshold, i.e. crosses one of two response boundaries. The upper response boundary was defined as selection of the optimal (80% reinforced) stimulus, whereas the lower response boundary was defined as selection of the suboptimal (20% reinforced) stimulus. RTs for choices of the suboptimal option where multiplied by −1 prior to model estimation, and we discarded for each participant the fastest 5% of trials in order to ensure that implausibly fast trials do not exert an undue influence on the modeled RT distributions. For comparison with the RLDDMs, we first examined a null model without learning (DDM_0_). Here, the RT on each trial *t* is distributed according to the Wiener First Passage Time (*wfpt*):

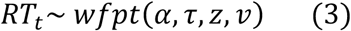

The boundary separation parameter *α* controls the speed-accuracy trade-off, such that smaller values of *α* lead to faster but less accurate responses. The drift rate *v* reflects the quality of the evidence, such that greater values of *v* give rise to more accurate and faster responses. Note that in the DDM_0_, *v* is constant and not affected by the learning process. The non-decision time *τ* models RT components related to motor and/or perceptual processing that are unrelated to the evidence accumulation process. The starting point parameter *z*, which we fixed at 0.5, models a bias towards one of the response boundaries.

Following earlier work^58,59,62,66^ we then incorporated the learning process (Equations 1 and 2) into the DDM by setting trial-wise drift rates to be proportional to the difference in Q-values between the optimal and suboptimal options via a simple linear linkage function:

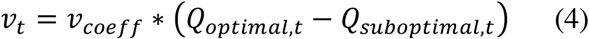

In the RLDDMs, the RT on trial *t* then depends on this trial-wise drift rate:

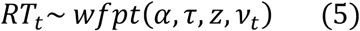

Here, *v_Coeff_* models the degree to which trial-wise drift rates scale with the value difference between options. In this model, increasing Q-value differences lead to both increased accuracy and faster RTs. Conversely, when Q-values are similar, choices will be both less accurate and slower.

Previous studies have also examined non-linear linkage functions^66–68^. However, in the present data, models with non-linear linkage functions failed to converge. We therefore focused on Eq. 4 originally proposed by Pedersen and colleagues^59^ and successfully applied in other learning tasks^62^. This simpler model nonetheless reproduced key patterns in the data, in particular the increase in accuracy over trials, the decrease in RTs over trials, as well as individual participant RT distributions.

In an earlier report^58^ we have additionally examined models in which *α* and/or *τ* were allowed to vary over the course of the experiment according to a power function. This modification of the RLDDM was motivated by the observation that, in gambling disorder participants, reinforcement learning only accounted for a relatively small part of the observed RT reductions over time, such that an additional mechanism was required to account for this observation. In contrast, in the present study, posterior predictive checks revealed that this was not the case in any drug condition. Therefore, we refrained from examining these more complex models here, instead focusing on models with constant *α* and *τ*.

### Hierarchical Bayesian models

Models were fit to all trials from all participants using a hierarchical Bayesian modeling approach with separate group-level Gaussian distributions for all parameters. We ran two types of models. First, we fit separate hierarchical models to the data from each drug condition and compared the model ranking across conditions. After confirming that the model ranking was unaffected by drug, we set up a combined hierarchical model in which parameters in the Placebo condition were modeled as the baseline, and deviations from placebo under L-dopa and Haloperidol were modeled for each parameter using additive shift parameters with Gaussian priors centered at 0. Posterior distributions were estimated using Markov Chain Monte Carlo via JAGS (Version 4.3)^69^ using the Wiener module^70^, in combination with Matlab (The MathWorks) and the *matjags* interface (https://github.com/msteyvers/matjags). For group-level parameter means in the placebo condition in the combined model, as well as in the separate models per drug condition, we used uniform priors defined over numerically plausible parameter ranges (see Supplemental Table S1). For drug-effects in the combined model we used Gaussian priors centered at zero (see Supplemental Table S1). For group-level standard deviations, we used uniform priors over numerically plausible ranges (see Supplemental Table S2).

For each model, we ran 2 chains with a burn-in period of 100k samples and thinning factor of 2. 10k additional samples were then retained for analysis. Chain convergence was assessed by examining the Gelman-Rubinstein convergence diagnostic 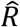, and values of 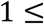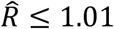 were considered as acceptable for all group-level and individual-subject parameters. Relative model comparison was performed via the estimated log pointwise predictive density (*elpd*)^71^, an approximation of the leave-one-out cross-validation accuracy of a model.

### Parameter recovery

We have previously reported parameter recovery analyses for RLDDMs in the context of the same task studied here^58^. These analyses revealed good parameter recovery for both individual-level and group-level parameters using the same Bayesian estimation methods applied here.

### Posterior predictive checks

To ensure that the best-fitting models captured key aspects of the data, in particular the increases in accuracy and reductions in RTs over the course of the experiment, we performed posterior predictive checks as follows. We simulated 10k data sets from each model’s posterior distribution, separately for each drug condition. For group-level posterior predicitive checks, per drug condition, we selected 1k simulated data sets, and for each simulated participant split trials into ten time bins. For each bin, we then calculated group average accuracy and RT as well as means and 2.5% and 97.5% percentiles over the simulated data. Simulated and data were then overlaid over the observed data. For individual subject posterior predictive checks, per drug condition, we again selected 1k simulated data sets. In first step, we simply overlaid observed and simulated RT distributions, separately for each condition. To examine how the models accounted for learning-related RT changes in individual subjects, we then split both simulated and observed trials into five time bins. For each bin, we calculated individual-subject means as well as means and 2.5% and 97.5% percentiles over the simulated data.

### FMRI data acquisition

MRI data were collected on a Siemens Trio 3T system using a 32-channel head coil. Each session, participants performed a single run of 60 trials (following a short break after completion of our previously reported task^19^). Each volume consisted of 40 slices (2 x 2 x 2 mm in-plane resolution and 1-mm gap, repetition time = 2.47s, echo time 26ms). We tilted volumes by 30° from the anterior and posterior commissures connection line to reduce signal drop out in the ventromedial prefrontal cortex and medial orbitofrontal cortex^72^. Participants viewed the screen via a head-coil mounted mirror, and logged their responses via the index and middle finger of their dominant hand using an MRI compatible button box. High-resolution T1 weighted structural images were obtained following completion of the cognitive tasks.

### FMRI preprocessing

All preprocessing and statistical analyses of the imaging data was performed using SPM12 (Wellcome Department of Cognitive Neurology, London, United Kingdom). Volumes were first realigned and unwarped to account for head movement and distortion during scanning. Second, slice time correction to the onset of the middle slice was performed to account for the shifted acquisition time of slices within a volume. Third, structural images were co-registered to the functional images. Finally, all images were smoothed (8mm FWHM) and normalized to MNI-space using the DARTEL tools included in SPM12 and the VBM8 template.

### FMRI statistical analysis

Error trials were defined as trials were no response was made within the 3sec response window, or trials that were excluded from the computational modeling during RT-based trial filtering (see above, for each participant, the fastest 5% of trials were excluded). We then set up first-level general linear models (GLMs) for each participant and drug condition. We used GLM1 for all main analyses, and GLM2 to reproduce a key analysis from Pessiglione et al.^17^

GLM1 included the following regressors:

1. onset of the decision option presentation
2. onset of the decision option presentation modulated by chosen – unchosen value
3. onset of the decision option presentation modulated by (chosen – unchosen value) squared
4. onset of the feedback presentation
5. onset of the feedback presentation modulated by model-based prediction error
6. onset of the decision option presentation for error trials and
7. onset of the feedback presentation for error trials.

To separate out effects of positive vs. negative prediction error coding, as done in Pessiglione et al.^17^, we set up a second first-level GLM.

GLM2 included the following regressors:

1. onset of the decision option presentation
2. onset of the decision option presentation modulated by chosen – unchosen value
3. onset of the decision option presentation modulated by (chosen – unchosen value) squared
4. onset of the feedback for positive prediction errors
5. onset of the feedback for negative prediction errors
6. onset of the decision option presentation for error trials and
7. onset of the feedback presentation for error trials.

To reproduce the analysis from Pessiglione et al.^17^, Figure 3, striatal activation peaks encoding model-based prediction errors were first identified using contrast 5 in GLM1 (see below for the metanalysis-based region of interest mask that was applied). From GLM2, we then extracted parameter estimates for positive and negative prediction errors at these peak voxels.

Values and prediction errors were calculated using the condition-specific group-mean learning rates of the best-fitting model RLDDM2 (see Table 1). All parametric regressors were z-scored within subject prior to entering the first level model^73^. Single-subject contrast estimates were then taken to a second-level random effects analysis using the flexible factorial model as implemented in SPM12 with a single within-subject factor of drug condition.

**Table 1.** Group-mean posterior positive (*η*_+_) and negative (*η*_−_) learning rates used for option value and prediction error calculation. These values were taken from RLDDM2 fitted separately to the data from each drug condition.

| Condition | $\eta_+$ | $\eta_-$ |
| --- | --- | --- |
| Placebo | .319 | .0004 |
| L-Dopa | .282 | .0149 |
| Haloperidol | .266 | .0001 |

We focused our analyses on reward-related circuits using a region of interest (ROI) mask provided by the Rangel Lab (https://www.rnl.caltech.edu/resources/index.html) which is based on two meta-analyses^74,75^. This mask was used for small-volume-correction for our analyses of chosen – unchosen value and prediction error, as well as for testing for drug effects. For all effects, we plot single-subject contrast estimates extracted from group-level activation peaks.

### Data and code availability

Raw trial-level and processed behavioral data are available on the OSF framework (*link will be made available upon publication*). Raw fMRI data cannot be shared becausee participants did not provide consent. However, group-level T- and F-maps for all main contrasts are available on OSF (*link will be made available upon publication*). Model code is likewise available on OSF (*link will be made available upon publication*).

## Results

### Model-agnostic analysis

Participants failed to respond within the allocated response time window only in very few trials (mean [range] number of misses Placebo: .61 [0 – 10], L-dopa: .61 [0 – 7], Haloperidol: .45 [0 – 4]). RT distributions per drug condition are shown in Figure 2a-c, with choices of the suboptimal option coded as negative RTs. A Bayesian signed rank test^64^ provided substantial evidence for performance above chance level under all drugs (see Figure 2d, all BF10>4000). Summary descriptive statistics for model-agnostic performance measures (accuracy, total rewards earned, median RTs) are shown in Figure 2d-f and Table 2, and results from Bayesian repeated measures ANOVAs^64^ are listed in Table 2. Numerically, both accuracy and median RTs were higher under Placebo compared to both L-dopa and Haloperidol, but Bayesian repeated measures ANOVAs provided in all cases moderate evidence in favor of the null hypothesis (Table 2).

**Figure 2.**
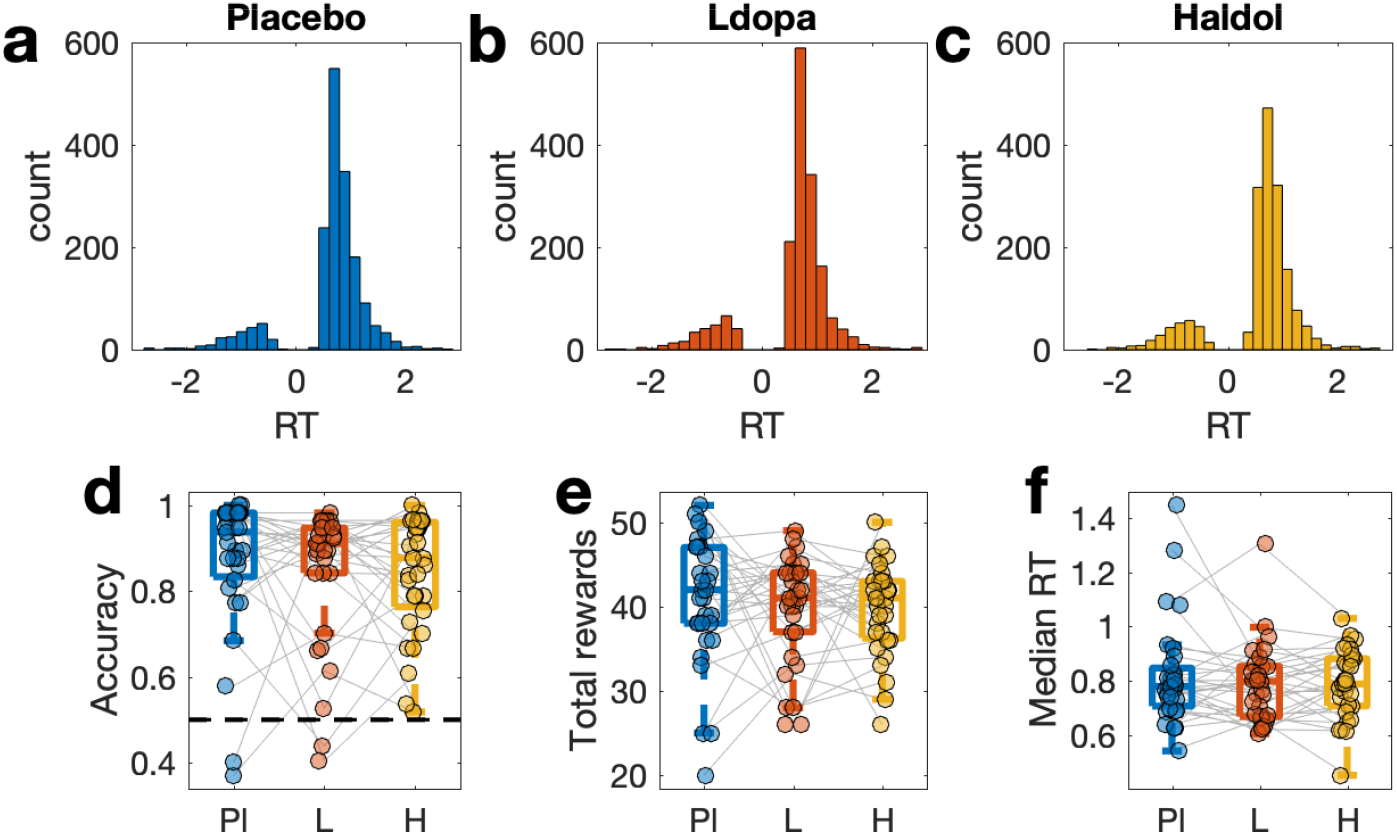
Response time (RT) distributions (n=31, within-subject design) under Placebo (a), L-dopa (b) and Haloperidol (c). Choices of the suboptimal options in (a-c) are coded as negative RTs, whereas choices of the optimal options are coded as positive RTs. d: Accuracy per drug condition (chance level is 0.5). e: Median RT per drug condition. Pl – Placebo, L – L-dopa, H – Haloperidol.

**Figure 3.**
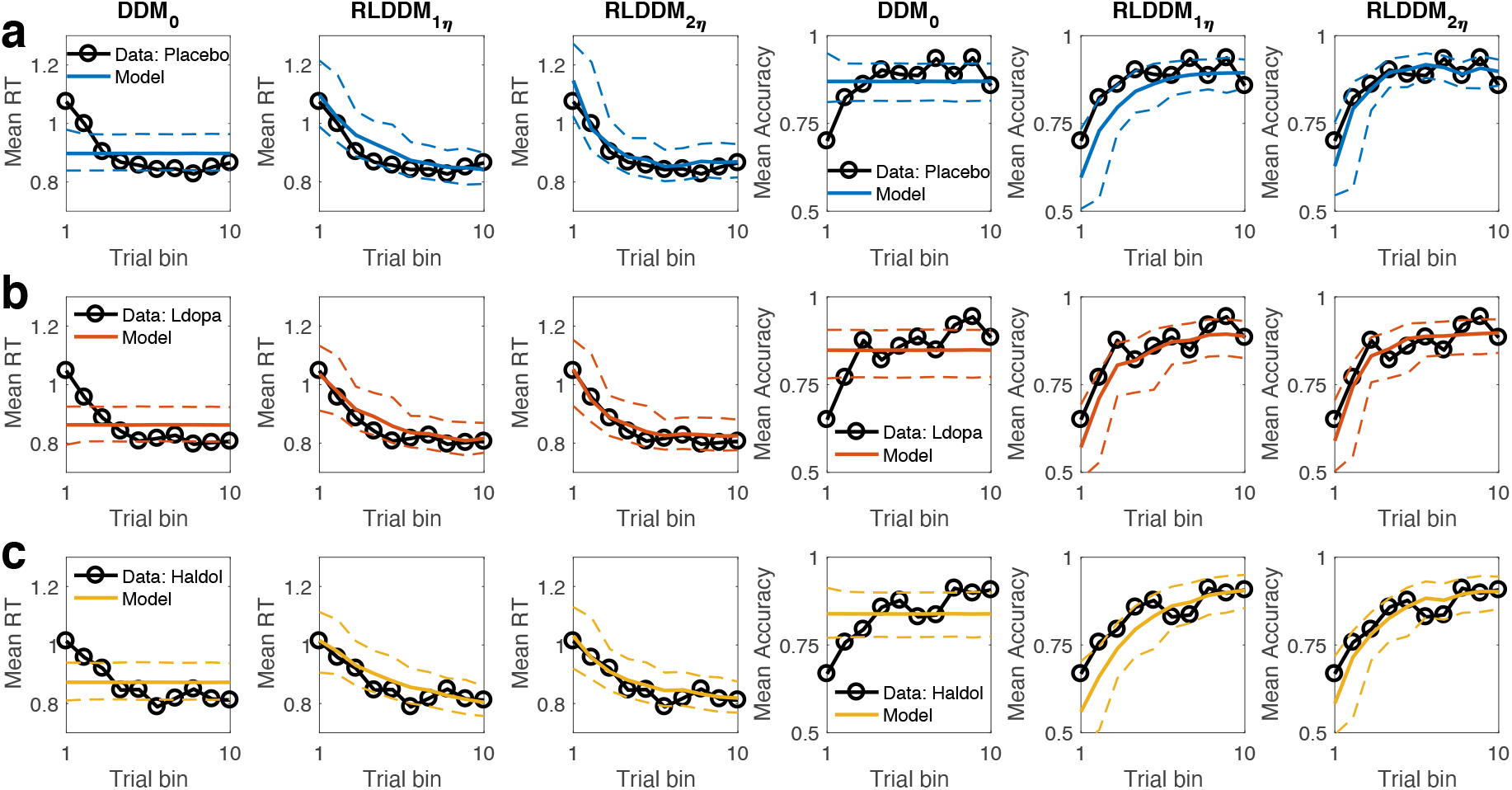
Group-level posterior predictive checks. DDM_0_: Null model without learning. RLDDM1η: Reinforcement learning DDM with a single learning rate. RLDDM2η: Reinforcement learning DDM with dual learning rates. Data and model simulations are shown for Placebo (a), L-dopa (b) and Haloperidol sessions (c). Left columns: observed RTs over time (black lines) and model predicted RTs (solid colored lines: means, dashed lines: +/− 95% percentiles). Right columns: observed accuracies over time (black lines) and model predicted accuracies (solid colored lines: means, dashed lines: +/− 95% percentiles).

**Table 2.** Descriptive statistics [Mean (STD)] for model-agnostic performance measures, and Bayes Factors in favor of the null model (BF01) from Bayesian repeated measures ANOVAs.

| Condition | Accuracy | Total rewards | Median RT |
| --- | --- | --- | --- |
| Placebo | .872 (.163) | 40.849 (7.781) | .820 (.188) |
| L-dopa | .842 (.161) | 39.710 (6.394) | .788 (.143) |
| Haloperidol | .840 (.136) | 39.645 (5.395) | .789 (.126) |
| *BF01* | 6.24 | 7.52 | 5.49 |

Given the within-subjects design, we also tested for a potential meta-learning effect across sessions. This analysis revealed moderate evidence in favour of the null model withouth a session effect (Supplemental Figure S1, BF01=8.41).

To directly replicate the analysis of the primary behavioral effect reported in Pessiglione et al.^17^ (difference in total rewards earned between L-dopa and Haloperidol conditions), we ran a frequentist paired t-test between L-Dopa and Haloperidol, which revealed no significant effect (t_30_=.045, p=.964). This was also the case when restricting the analysis to only those subjects who received L-Dopa and Haloperidol on the first session (t_19_=−.943, p=.358).

### Model comparison

We next compared three computational models (see methods section). As a reference, we first fit a null model (DDM_0_) without a learning component (i.e. constant drift rate across trials). We then examined two reinforcement learning drift diffusion models (RLDDMs) that included a linear mapping from Q-value differences to trial-wise drift rates^59,60,62^ (see Eq. 5) with either a single learning rate (RLDDM1) or dual learning rates η for positive vs. negative prediction errors (RLDDM2).

Model comparison was performed using the estimated log pointwise predictive density (-elpd)^71^. The RLDDM with dual learning rates outperformed both the single learning rate RLDDM and the DDM_0_ model without learning (see Table 3). This model ranking replicated across all three drug conditions.

**Table 3.** Model comparison separately per drug condition. We examined reinforcement learning drift diffusion models (RLDDMs) with a single learning rate η (RLDDM 1) vs. separate learning rates for positive and negative prediction errors (RLDDM 2). We also included a null model without learning (DDM_0_). Model comparison was conducted using the estimated log pointwise predictive density (-elpd)^71^ where smaller values indicate a better fit.

| Model | $\eta$ | Placebo | | L-dopa | | Haloperidol | |
| --- | --- | --- | --- | --- | --- | --- | --- |
|  |  | -elpd (SE) | Rank | -elpd (SE) | Rank | -elpd (SE) | Rank |
| $DDM_0$ | - | 296.5 (52.1) | 3 | 375.9 (52.8) | 3 | 480.2 (51.2) | 3 |
| RLDDM 1 | Single | 178.7 (47.7) | 2 | 271.6 (50.0) | 2 | 412.7 (49.5) | 2 |
| RLDDM 2 | Dual | 67.3 (48.5) | 1 | 195.4 (51.2) | 1 | 336.8 (50.7) | 1 |

### Posterior predictive checks

We next ran posterior predictive checks to examine the degree to which the best-fitting model (RLDDM2) accounted for key patterns in the data, in particular the increase in accuracy and the reduction in RTs over the course of learning. We simulated 10k datasets from each model’s posterior distribution. We then binned trials into ten bins over the course of learning, and computed mean accuracies and RTs per bin for both the observed data and across a 1k subset of simulated data sets, separately for each drug condition. Checks are depicted in Figure 3.

Since the DDM_0_ predicts constant accuracies and RTs over trials, it cannot reproduce the observed increases in accuracy and reductions in RTs over the course of learning (Figure 3). In contrast, both RLDDMs reproduced the learning-related reductions in RTs (Figure 3 left) and the increases in accuracy (Figure 3, right). However, RLDDM1 tended to underestimate both effects. In particular, it underpredicted accuracies (Figure 3, right), and somewhat overpredicted RTs (Figure 3, left). RLDDM2 provided a better account of these effects, in particular with respect to RTs. In line with the model comparison, this pattern was observed in all drug conditions.

It was recently suggested that RLDDMs might fail to account for the full RT distributions^76^. We therefore ran additional posterior predictive checks that examined predictions of 17^th^, 50^th^ and 83^rd^ percentiles of the RT distributions (Supplemental Figure S1). Again, RLDDM2 reproduced these data well. Finally, RLDDM2 also reproduced individual subject RT distributions (Supplemental Figures S2–S4), and accounted for the evolution of RTs in individual subjects over the course of learning (Supplemental Figure S5–S7), where for all subjects the observed data fell within the 95% prediction interval of the simulated RTs.

### Analysis of drug effects on model parameters

We next analyzed the posterior distributions of the RLDDM2, focusing on the combined model. Recall that here, the Placebo condition was modeled as the baseline, and drug effects were modeled as additive changes from baseline, for each parameter. Figure 4 (top row) depicts the posterior distributions of the group mean of each parameter under Placebo, whereas the mid- and center rows show effects of L-dopa and Haloperidol on each parameter (see also Table 3). Drug effects were quantified in three ways. First, we examined the overlap of the 95% HDI of each condition effect with zero. Only for drug effects on the boundary separation parameter did zero fall outside of the 95% HDI (Figure 4a). We next computed regions-of-practical equivalence^77^ (ROPEs) in terms of ± 0.1 SD of the drug effects. Of all parameters, only the 95% HDI for the boundary separation effects fell beyond the ROPEs (Figure 4a). We finally computed directional Bayes Factors (Table 3) quantifying the relative evidence for increases vs. decreases in a parameter as a function of the different drug conditions. Decreases in boundary separation under both L-dopa and Haloperidol were substantially more likely, given the data, than increases (Table 4).

**Figure 4.**
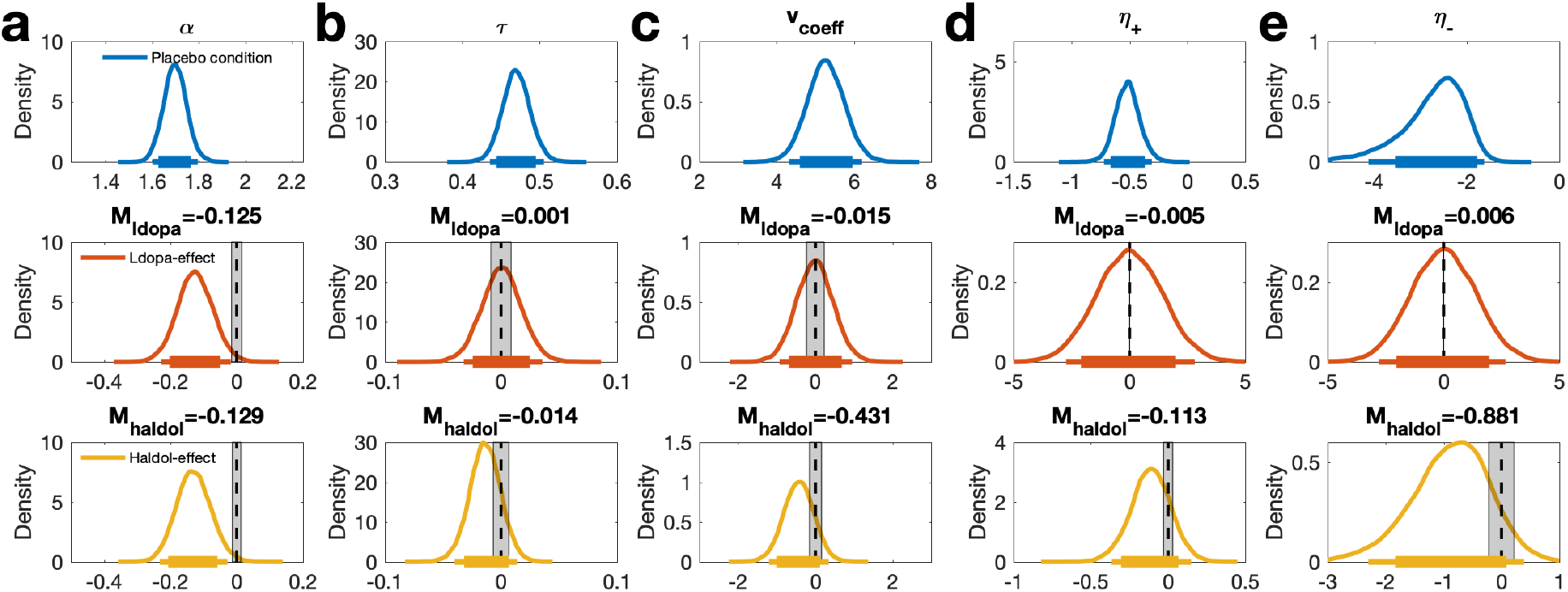
Drug effects on RLDDM parameters. a: boundary separation (*α*), b: non-decision time (*τ*), c: value coefficient of the drift rate (*ν*_coeff_), d: positive learning rate (η_+_) in standard normal space, e: negative learning rate (η_−_) in standard normal space. Top row: Posterior distributions for each parameter under Placebo. Center row: Posterior distributions of L-dopa-effects on each parameter (M_ldopa_ refers to the mean). Bottom row: Posterior distributions of Haloperidol-effects on each parameter (M_haldol_ refers to the mean). Solid (thin) horizontal lines denote 85% (95%) highest posterior densities. Shaded areas denote Regions of Practical Equivalence (ROPEs)^77^ ± 0.1 SD.

**Table 4.**
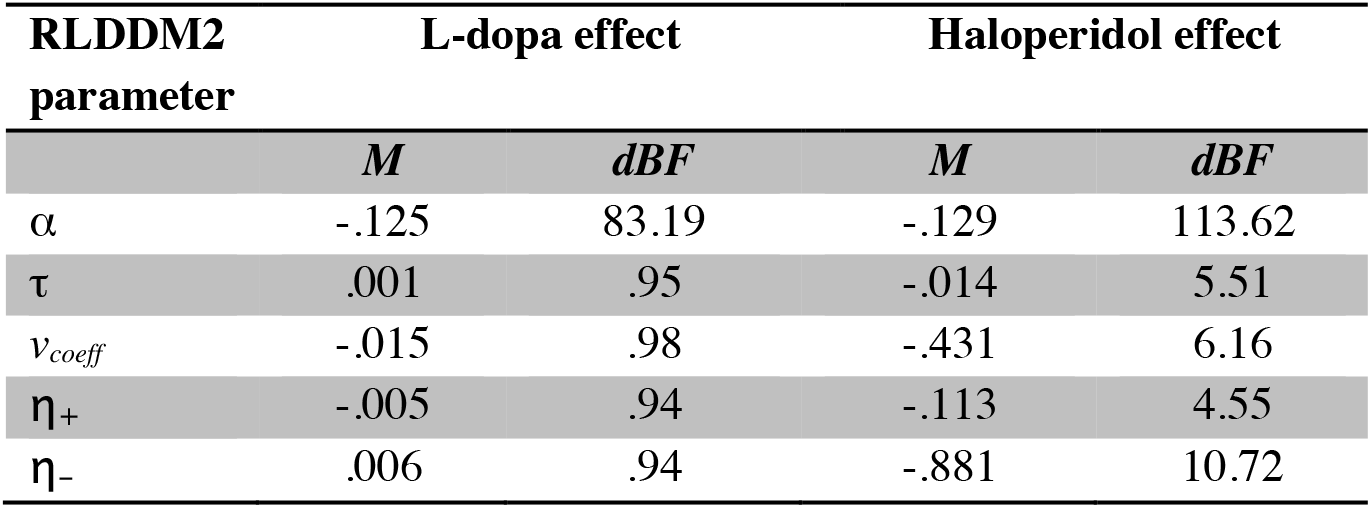
Drug effects on model parameters: Mean posterior drug effects on model parameters (M_diff_) relative to the Placebo condition, for L-dopa (left) and Haloperidol (right), as well as Bayes factors testing for directional effects (dBF). dBF values > 1 quantify the degree of evidence for a reduction in a parameter compared to placebo, compared to the evidence for an increase. dBF values < 1 reflect the reverse.

| RLDDM2<br>parameter | L-dopa effect |  | Haloperidol effect |  |
| --- | --- | --- | --- | --- |
| | $M$ | dBF | $M$ | dBF |
| $\alpha$ | -.125 | 83.19 | -.129 | 113.62 |
| $\tau$ | .001 | .95 | -.014 | 5.51 |
| $\nu_{\text{coeff}}$ | -.015 | .98 | -.431 | 6.16 |
| $\eta_+$ | -.005 | .94 | -.113 | 4.55 |
| $\eta_-$ | .006 | .94 | -.881 | 10.72 |

We repeated the analysis of drug effects using a modeling scheme in which separate models were fit to the data from each drug condition. This reproduced the effects observed in the combined model (Supplemental Figure S9, Supplemental Table S3). Again, the only parameter showing drug effects was the boundary separation, which was reduced in both drug conditions, compared to Placebo.

Finally, to link the modeling results back to individual behavior, we examined the degree to which individual differences in drug effects on boundary separation accounted for differences in RTs between conditions. Across participants, greater RT differences between Placebo and drug were associated with a greater reduction in boundary separation (Figure 5a, L-dopa: *r*=−.48, BF10=8.173, Figure 5b, Haloperidol: *r*=−.728, BF10=6557) - decision threshold reductions under both L-dopa and Haloperidol were associated with corresponding RT reductions.

**Figure 5.**
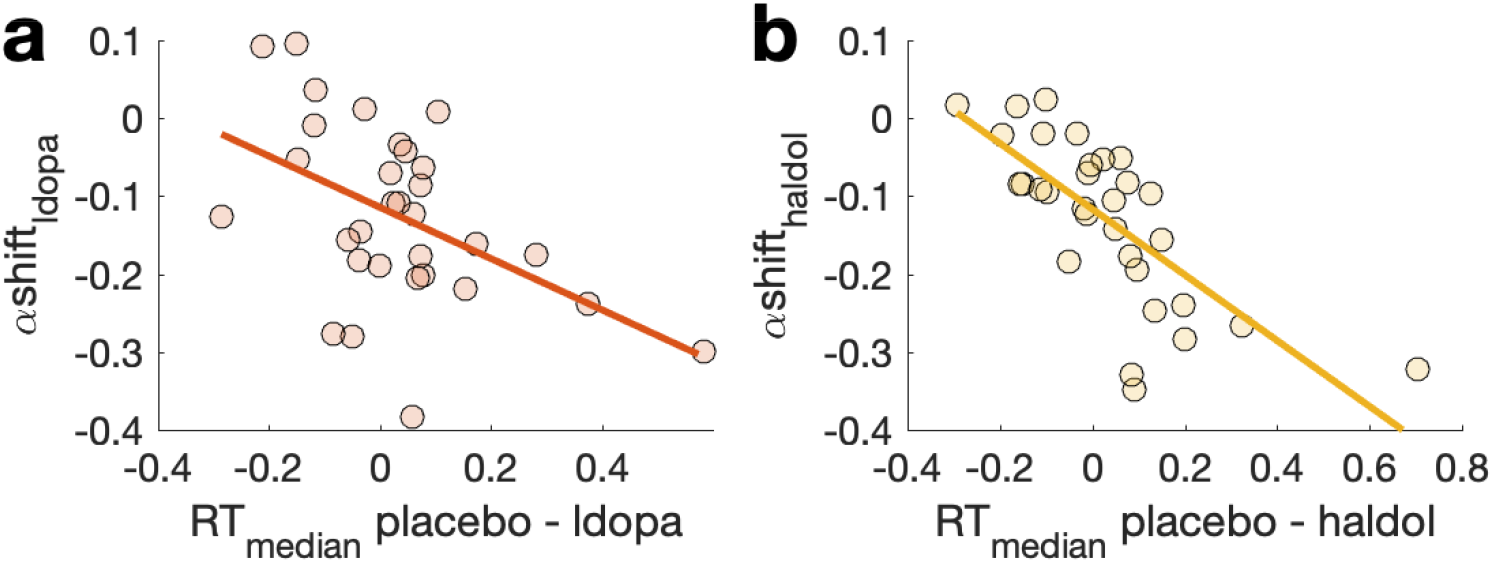
Individual differences in drug effects on boundary separation α from the combined RLDDM2 (y-axis), were associated with RT differences between placebo and drug (differences in median RTs, x-axis) both for L-dopa (a, *r*=−.48, BF10=8.173) and Haloperidol (b, *r*=−.728, BF10=6557).

### FMRI results

FMRI analyses focused on a single *a priori* region of interest (ROI) based on two meta-analyses of value effects (see methods, including ventral striatum, ventromedial prefrontal cortex, posterior cingulate and anterior cingulate). Using parametric measures derived from our computational model (chosen – unchosen value, model-based prediction error), we first aimed to replicate effects of model-based chosen – unchosen value in vmPFC/mOFC and prediction error in ventral striatum. Main effects across drug conditions in our reward ROI (see methods section) showed that both effects replicated (Table 5).

**Table 5.** Replication analysis for three model-based measures (main effect across drug conditions). Small volume correction (SVC) used an a priori region of interest mask across two meta-analyses of reward value effects^74,75^ (see methods section).

| Contrast / Region | Coordinates | Peak T-value | $p(\text{FWE})_{\text{SVC}}$ |
| --- | --- | --- | --- |
| Chosen-unchosen value |  |  |  |
| vmPFC / mOPFC | 6 24 -12 | 4.11 | .032 |
| Reward prediction error |  |  |  |
| Left ventral striatum | -9 9 -9 | 5.95 | <.001 |
| Right ventral striatum | 12 9 -10 | 4.93 | .003 |
| vmPFC / mOFC | -6 32 -15 | 4.19 | .026 |
| vmPFC | -6 54 3 | 4.08 | .036 |

Chosen value effects are shown in Figure 6a, and prediction error effects are shown in Figure 6b. In a next step, we tested for drug effects on the same three effects via F-contrasts testing for main effects of drug. In none of the contrasts did we observe any effects that survived correction for multiple comparisons across the reward ROI. We also did not observe drug effects when running an FWE-corrected whole-brain analysis on these three effects. Finally, we tested for drug effects on stimulus-onset and feedback-onset related effects, again using whole-brain FWE correction. No significant effects were observed.

**Figure 6.**
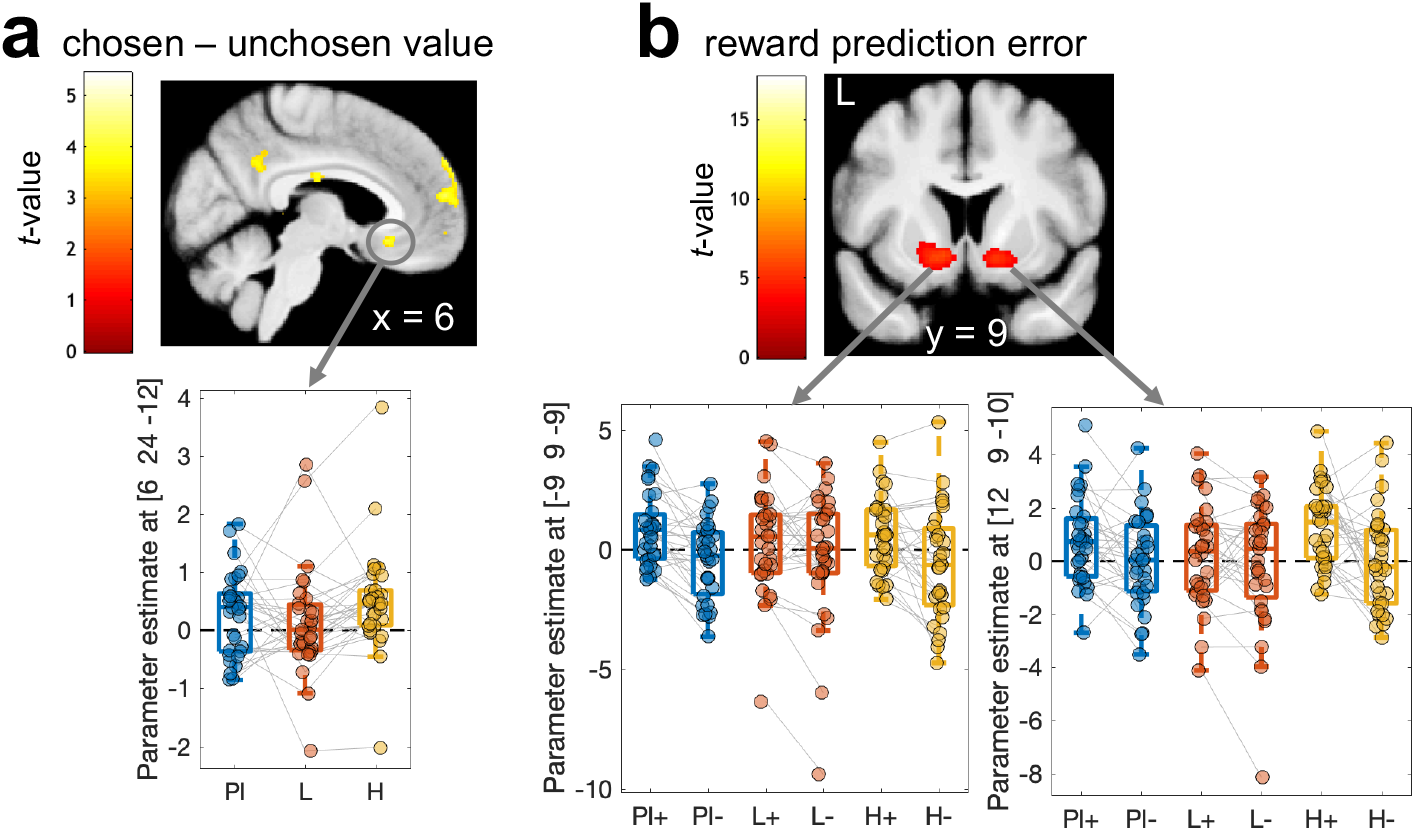
FMRI results for model-based chosen – unchosen value (a) and reward prediction error (b). Correction for multiple comparisons was performed using a meta-analysis-based region-of-interest mask (see Table 5 and methods section). a: Chosen – unchosen value was represented in mOFC/vmPFC. b: We first identified striatal regions coding for model-based prediction errors by computing the main effect of prediction error coding across all drug conditions using GLM1. To reproduce the analysis of Pessiglione et al.^17^, we then extracted parameter estimates at striatal peak voxels (see Table 5) from GLM2 to obtain separate parameter estimates for positive (+) and negative (−) prediction errors. Maps are thresholded at *p*<.001 *uncorrected* for display purposes, and projected on the group mean T1 scan. Pl – Placebo, L – Levodopa, H – Haloperidol.

In a last step, we reproduced the analysis from Pessiglione et al.^17^ and separated out positive and negative prediction error effects in bilateral ventral striatal regions that encoded model-based prediction errors (see Figure 6b and Table 5) in GLM1. Using a separate GLM that included separate predictors for positive and negative prediction errors, the corresponding parameter estimates were extracted from bilateral ventral striatum (Figure 6b). While Pessiglione et al.^17^ reported a greater contrast between positive and negative striatal prediction errors under L-dopa compared to Haloperidol, this was not the case in our data (Figure 6b). Bayesian ANOVAs with the factors prediction error (positive vs. negative) and drug (Placebo vs. L-dopa vs. Haloperidol) revealed strong (left ventral striatum, BF=20.49) and moderate (right ventral striatum, BF=6.42) evidence in favor of a model that only included a main effect of prediction error, but no drug effect or drug*prediction error interaction.

## Discussion

We used a stationary reinforcement learning task^17^ in combination with a pharmacological manipulation of dopamine (DA) neurotransmission (Placebo, 150mg L-dopa, 2mg Haloperidol) and fMRI to address two core research questions. First, we aimed to replicate the previously reported beneficial effect of L-dopa (vs. Haloperidol) on reinforcement learning and striatal prediction error signaling^17^. This replication was not successful - there was no evidence for improved learning under L-dopa. In contrast, Bayesian analyses provided moderate evidence in favor of the null model. Perhaps unsurprisingly, given the lack of a behavioral effect, we also did not observe drug effects on positive vs. negative prediction error coding in ventral striatum, again contrasting with Pessiglione et al.^17^, but in line with recent studies using different reinforcement learning tasks^19,20^. Potential reasons for these unsuccessful replications are discussed further below. Second, we leveraged recently developed combined reinforcement learning drift diffusion models (RLDDMs)^59–63^ to directly test a recently proposed computational account of dopamine in regulating decision thresholds during action selection^38^. In line with this account, computational modeling revealed reduced decision thresholds under both L-dopa and Haloperidol, compared to Placebo. The latter effect is consistent with the present Haloperidol dosage of 2mg increasing (rather than decreasing) striatal DA by blocking presynaptic autoreceptors (see discussion below).

We aimed to replicate the core behavioral finding from Pessiglione et al.^17^, but for practical reasons deviated from their experimental design in a number of ways. First, the drug dosages in the two projects differed slightly – we used 150mg of L-dopa (compared to 100mg in Pessiglione et al.) and 2mg of Haloperidol (compared to 1mg in Pessiglione et al.). The L-Dopa dosage was chosen to keep drug dosages constant across multiple pharmacological studies in a larger project, in which related processes^78,79^ where investigated. The Haloperidol dosage was selected to keep drug dosages comparable to planned studies investigating other effects^80,81^, and prior to completion of our human work suggesting that 2mg Haloperidol might elicit effects more compatible with an increase in DA transmission^32,67^. Second, we only included the gain condition, because here, the primary behavioral effect was observed. Although we doubled the number of gain trials (i.e. we used two pairs of symbols), our task version was likely still easier than the one used by Pessiglione et al.^17^, who used three pairs of symbols. Furthermore, this isolation of the gain condition might have affected drug effects on learning. The reason is that, in our task version, the mean initial reward expectation was positive (there were only gains or reward omissions), yielding both positive and negative prediction errors during initial learning. In contrast, in Pessiglione et al., the mean initial reward expectation was zero (there were equally many gain and loss trials), yielding only positive Prediction errors in the gain condition during initial learning. This might have masked drug effects. Third, we increased the sample size to n=31, and applied a within-subjects design as opposed to a between-subjects design. This might have induced learning across sessions, which could have masked drug effects. However, in contrast with this idea, we observed no performance changes over time, and focusing the analysis only on the first session likewise yielded no drug effects. The most straightforward replication attempt of the behavioral effect, a comparison of total rewards obtained between the L-dopa and Haloperidol conditions via a frequentist paired t-test, yielded no significant effect. Bayesian analyses, in contrast, yielded moderate evidence in favor of the null hypothesis.

We consider the isolation of the gain condition to be the most likely reason for the lack of replication, although other accounts are possible. For example, differences in dosages migh account for the lack of a drug effect on performance, but we consider this unlikely, for several reasons. First, 150mg of L-dopa have yielded positive behavioral effects in a range of other studies^15^. Second, 2mg of Haloperidol is a dosage that likely predominantly affects presynaptic autoreceptors and thus likely increases (rather than decreases) striatal DA release^8,27–32,67^. Note that this generally questions a common interpretation of a downregulation of striatal DA using even lower dosages of Haloperidol^17,78^. Nonetheless, even if one would argue in favor of a DA-downregulation account of 1mg or 2mg of haloperidol (which we consider unlikely), in this case one would if anything expect more pronounced effects of our 2mg dosage compared to Pessiglione et al.’s 1mg dosage. Yet, inconsistent with this idea, no effects were observed. Also, the relatively small sample size of the original study might have led to an increased risk of false positives. Finally, there are substantial individual differences with respect to pharmacological effects of DA drugs^1^ and it is possible that such individual differences might have also contributed to the inconsistencies between studies.

Extending previous work, we applied RLDDMs to directly examine effects of DA on the dynamics underlying action selection. Model comparison revealed that the data were best accounted for by an RLDDM with separate learning rates for positive and negative prediction errors (RLDDM2), and the model ranking was replicated in each drug condition. We have previously confirmed good parameter recovery for RLDDMs in this task^58^. Posterior predictive checks likewise confirmed that RLDDM2 provided a good account of learning-related increases in accuracy and decreases in RT. Likewise, additional checks across different percentiles of the RT distributions revealed that RLDDM2 provided a good account of the data (Supplemental Figure S1). Although RLDDMs have been suggested to provide a comparatively poor account of full RT distributions^76^, this was not the observed here. Finally, we confirmed that RLDDM2 also provided a good account of individual-subject RT distributions in each drug condition (Supplemental Figures S3–S5), consistent with prior work^67,82^. It also reproduced learning-related reductions in RTs over the course of learning (Supplemental Figures S6–S8). In contrast to earlier work^54,61,63,67,68^, in the present data, models with a non-linear mapping from value differences to trial-wise drift rates failed to converge (potentially due to the lower number of trials compared to earlier work), and we therefore focused on the simpler model with a linear linkage function^59,62^.

RLDDM2 revealed that, compared to Placebo, both L-dopa and Haloperidol reduced decision-thresholds. This effect was observed both in the combined model across all drug conditions (Figure 4) and in a control analysis using separate models for each drug condition (Supplemental Figure S9, Supplemental Table S3). Although we did not observe evidence for drug effects on model-agnostic behavioral measures, the overall pattern of behavioral results is nonetheless consistent with the observed reduction in decision thresholds: numerically, under both L-dopa and Haloperidol, accuracy was lower, and median RTs were faster. Furthermore, individual differences in drug-induced decision threshold reductions accounted for individual differences in RT differences between conditions. This argues against the idea that e.g. excessive shrinkage of parameters modeling drug effects might have driven drug effects in the combined RLDDM.

We observed similar reductions in decision thresholds following L-dopa and Haloperidol administration, two very different dopaminergic agents. By increasing substrate availability, L-dopa is assumed to generally increases DA availability. In contrast, Haloperidol is a D2 receptor antagonist that likely exhibits dose-dependent effects on striatal DA release. Specifically, lower dosages are thought to predominantly affect presynaptic inhibitory autoreceptors^26^, thereby increasing in DA release^8,27–32^. Along similar lines, other D2 antagonists potentiate striatal effects at lower dosages^24^, and attenuate them at higher dosages^33^. A Haloperidol dosage of 2mg has been shown to substantially upregulate human striatal responses^32^. Notably, however, the dose-dependency of Haloperidol might additionally be region-dependent^23^, which further complicates interpretation of the effects.

Our finding of reduced thresholds following pharmacological increases in DA also resonate with some findings in Parkinson’s Disease (PD). PD patients when tested ON vs. OFF their DA medication sometimes show increased speed but impaired accuracy^83^, and a reduced ability to suppress premature actions^84^, consistent with reduced decision thresholds. However, other studies did not find evidence for reduced decision thresholds in PD patients ON vs. OFF medication during perceptual decision-making^85^.

This null-effect in perceptual decision-making in PD patients might also point to a more general pattern – the specific domain of the decision problem might determine the degree to which DA contributes to threshold adjustments. During perceptual decision-making, Bromocriptine, a DA agonist, did likewise not modulate decision thresholds^56^. As noted by the authors, this might also be due to dose-dependent presynaptic effects of Bromocriptine, which might have resulted in a net reduction in DA transmission^56^. But another possibility is that DA might specifically contribute to threshold adjustments in value-based decision settings^54^. This idea resonates with the role of dopamine in regulating response vigor^34–37^. In some of these accounts, DA is thought to signal whether increases in cognitive or physical effort (in some cases equivalent to increases in response rate) are worthwhile^37,42,86,87^. Adjustments in decision thresholds would constitute a simple computational mechanism to accomplish this. Reductions in decision thresholds during decision-making under high DA and increases in response vigor in effort-related tasks might thus both serve the same purpose of (potentially) increasing the reward rate. Such an account would predict that action selection in the context of high reward options (where the cues themselves likely elicit phasic DA release during the choice phase^13^) would likewise lead to a downregulation of decision thresholds. Notably, this is exactly what has been observed in recent work^63,88^.

Increased DA is thought to shift the activation balance between striatal *go* and *no go* parthways towards the *go* pathway^13,14,39^, thereby fascilitating action execution vs. inhibition. In some models, separate *go* and *no go* action weights are modeled for each action^13,14^. An increase in DA during choice (be it pharmacological, as in the present study, or incentive-based^63,88^) would then for each action boost the contrast of *go* and *no go* action weights^13,14^, leading to a general increase in the probability of action initiation. In a sequential sampling modeling framework, this would be captured by a reduction of decision thresholds (reduced boundary separation). Conceptually, this is related to the idea that DA signals the (subjective) precision of beliefs^40–42^. If one’s belief in the precision of action weights is increased under high DA, this would naturally lead to accepting less evidence prior to committing to a decision. However, given the lack of drug effects on fMRI results during both choice and feedback phases of the present task, our imaging results remain agnostic with respect to a modulation of striatal computations. Furthermore, given the different mechanisms of action, it remains a possibility that both drugs might have affected decision thresholds via different routes.

The present study has a number of limitations that need to be acknowledged. First, we only tested male participants, limiting the generalizability of our findings. Second, the present sample size of n=31 is likely still too low to comprehensively examine potential non-linear baseline-dependent drug effects^1,19,89^. Third, the task was performed following completion of a separate learning task^19^, i.e. about 60min post ingestion of L-dopa. L-dopa reaches peak plasma levels around 30-60 min post ingestion, with a plasma half-life of about 60-90 min^90,91^. The present task was therefore likely performed past the time point of peak L-dopa plasma levels, but likely before the plasma half-life was reached. Although it is possible that this timing may have contributed to the lack of L-dopa effects on learning and neural prediction error signaling, we consider this unlikely, for several reasons. First, in the learning task performed directly prior to the task reported here^19^, as well as in other studies^20^, L-dopa likewise failed to modulate striatal prediction error signaling. Second, the robust L-dopa effect on boundary separation argues against a drug timing account to explain the lack of modulation of the prediction error response.

To conclude, we failed to replicate the beneficial effect of L-dopa (vs. Haloperidol) on reinforcement learning in a reward context, as well as the proposed mechanistic account of an enhanced striatal prediction error response mediating this effect. Bayesian analyses in all cases provided at least moderate evidence in favor of the null hypothesis. Differences in experimental design between studies likely account for this lack of replication. In contrast, diffusion modeling revealed robust effects of both L-dopa and Haloperidol on decision thresholds. This provides causal evidence for a recently proposed computational account of the role of DA in action selection^38,39^, and is consistent with both drugs boosting action-specific activation contrasts between striatal *go* and *no go* pathways during choice^13,14^. Such a threshold modulation account of DA can potentially bridge circuit-level accounts of action selection in the basal ganglia^13,39^ with a proposed role of dopamine in regulating response vigour^34–37^.

## Acknowledgements

This work was funded by Deutsche Forschungsgemeinschaft (PE 1627/5-1 to J.P.). We thank the MRI staff at the Institute for Systems Neuroscience, UKE, Hamburg, for support during data acquitision, and Mathias Pessiglione for helpful discussions and comments on an earlier version of the manuscript.

## Author contributions

J.P., K.C. and A. W. designed the study. K.C. and F.G. performed the study. J.P., A.W. and B. W. analyzed the data. J.P. wrote the paper and all authors provided revisions. J.P. supervised the project.

## Supplemental material

**Supplemental Table S1.** Overview of priors for group-level means.

| Parameter |  | Mean<br>(placebo condition) | Mean<br>(drug effects) |
| --- | --- | --- | --- |
| $\alpha$ | Boundary separation | $U(.01, 5)$ | $\mathcal{N}(0, 2)$ |
| $\tau$ | Non-decision time | $U(.1, 2)$ | $\mathcal{N}(0, 2)$ |
| $v_{coeff}$ | Drift rate coeff | $U(-100, 100)$ | $\mathcal{N}(0, 2)$ |
| $\eta_+, \eta_-$ | Learning rates | $U(-5, 5)$ | $\mathcal{N}(0, 2)$ |

**Supplemental Table S2.**
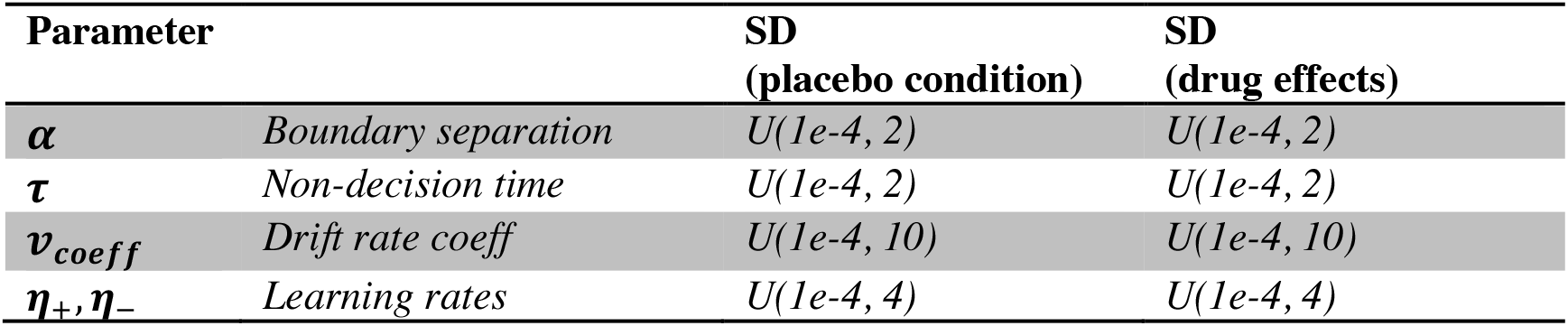
Overview of priors for group-level standard deviations.

| Parameter |  | SD<br>(placebo condition) | SD<br>(drug effects) |
| --- | --- | --- | --- |
| $\alpha$ | Boundary separation | $U(1e-4, 2)$ | $U(1e-4, 2)$ |
| $\tau$ | Non-decision time | $U(1e-4, 2)$ | $U(1e-4, 2)$ |
| $v_{coeff}$ | Drift rate coeff | $U(1e-4, 10)$ | $U(1e-4, 10)$ |
| $\eta_+, \eta_-$ | Learning rates | $U(1e-4, 4)$ | $U(1e-4, 4)$ |

**Supplemental Figure S1.**
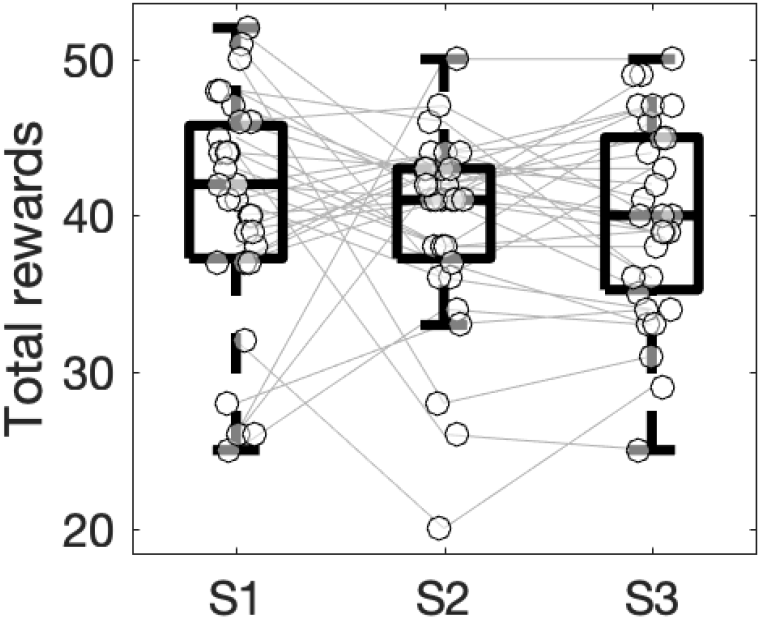
Total rewards earned per session. Bayesian repeated measures ANOVA yielded moderate evidence against an effect of session (BF01=8.41).

**Supplemental Figure S2.**
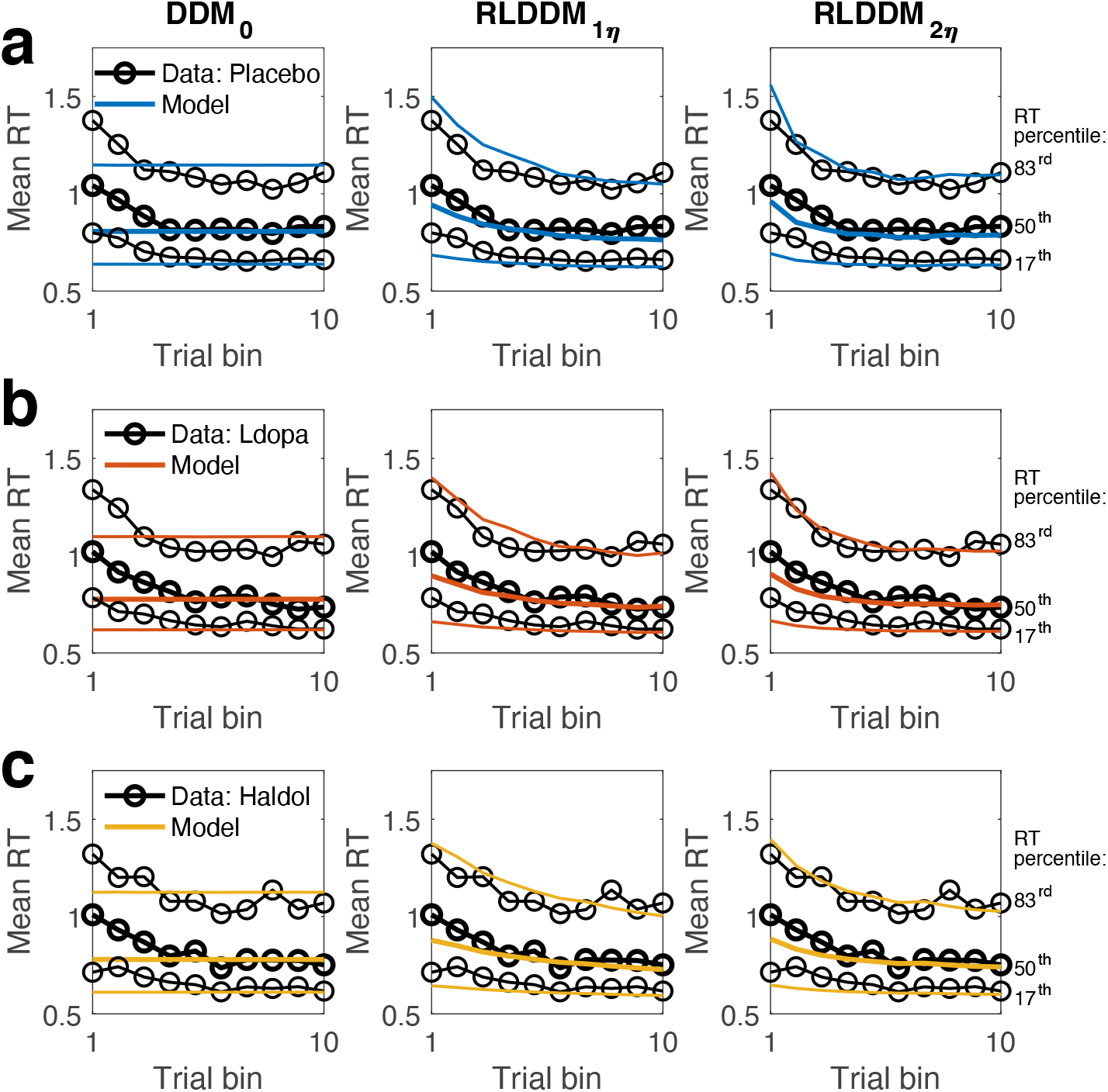
Posterior predictive checks for RTs using RLDDM2 (a: Placebo; b: ldopa; c: Haldol). Black lines denote mean RTs across subjects per trial bin for median RT (solid black line) and 83^rd^ and 17^th^ percentiles (think black lines). The colored lines show the respective simulations based on the posterior distributions.

**Supplemental Figure S3.**
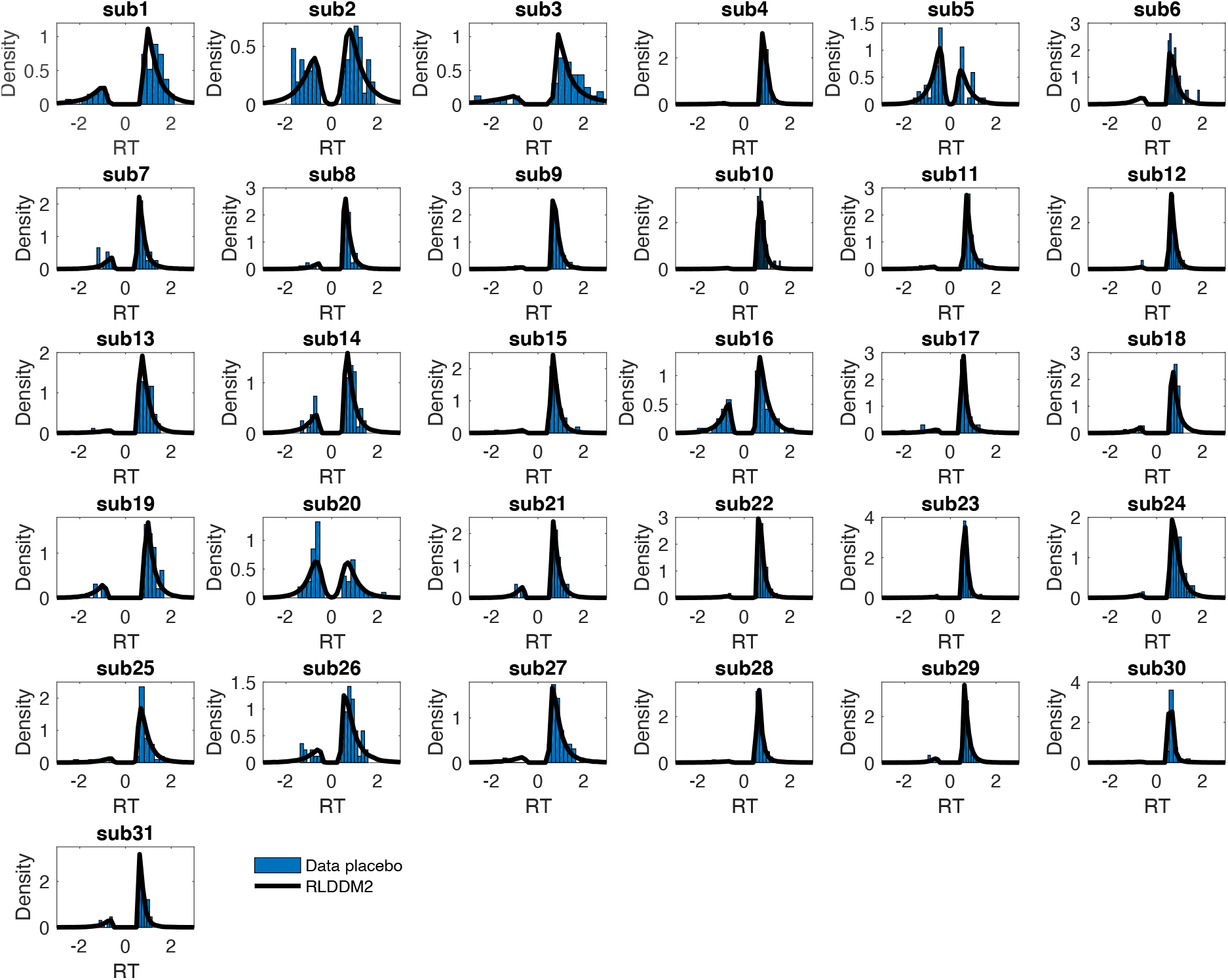
Single-Subject posterior predictive checks for the placebo condition. Histograms depict single-subject RT distributions (suboptimal choices are plotted as negative RTs). Solid black lines show smoothed histograms across 1k datasets simulated from the RLDDM2 posterior distribution.

**Supplemental Figure S4.**
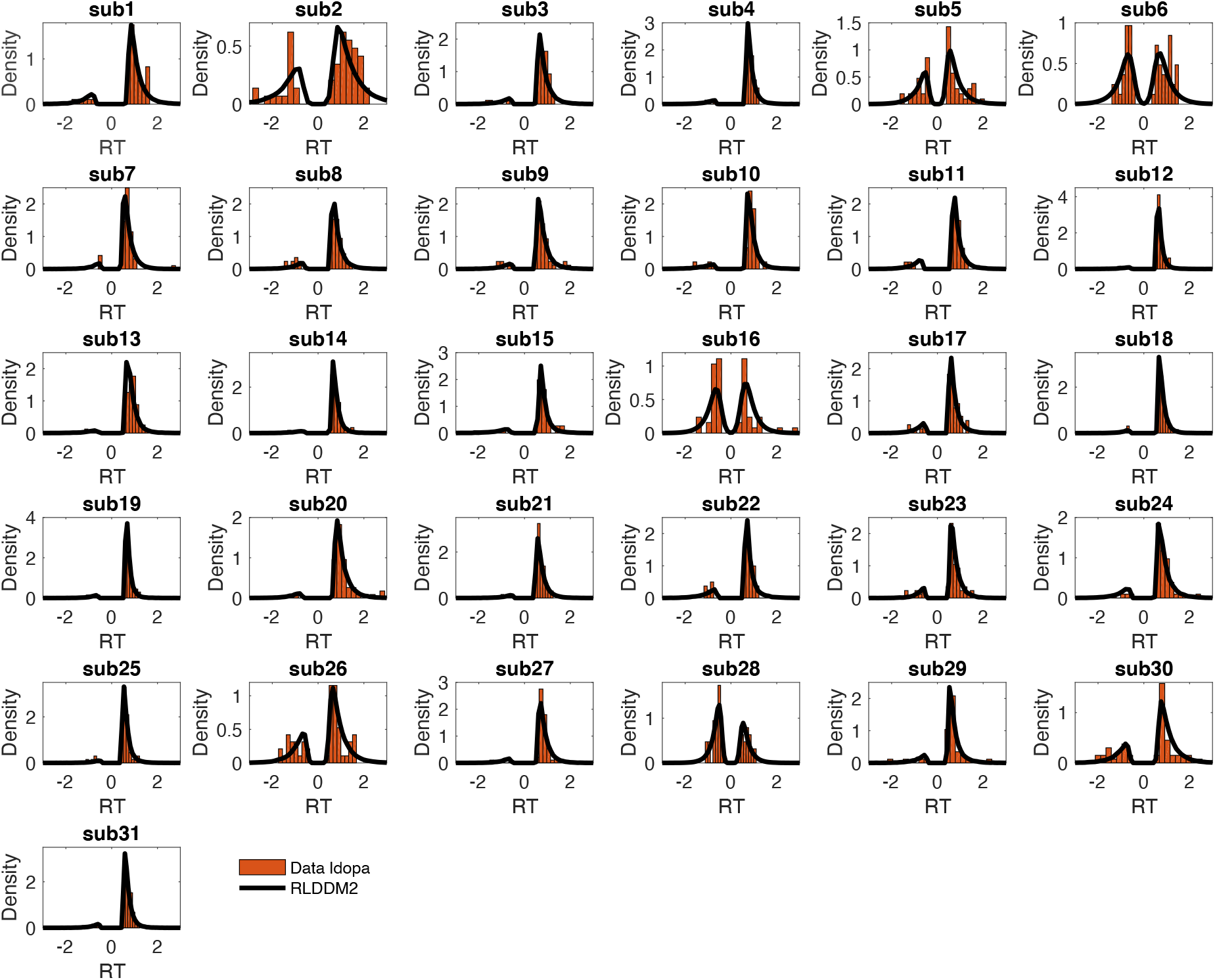
Single-Subject posterior predictive checks for the ldopa condition. Histograms depict single-subject RT distributions (suboptimal choices are plotted as negative RTs). Solid black lines show smoothed histograms across 1k datasets simulated from the RLDDM2 posterior distribution.

**Supplemental Figure S5.**
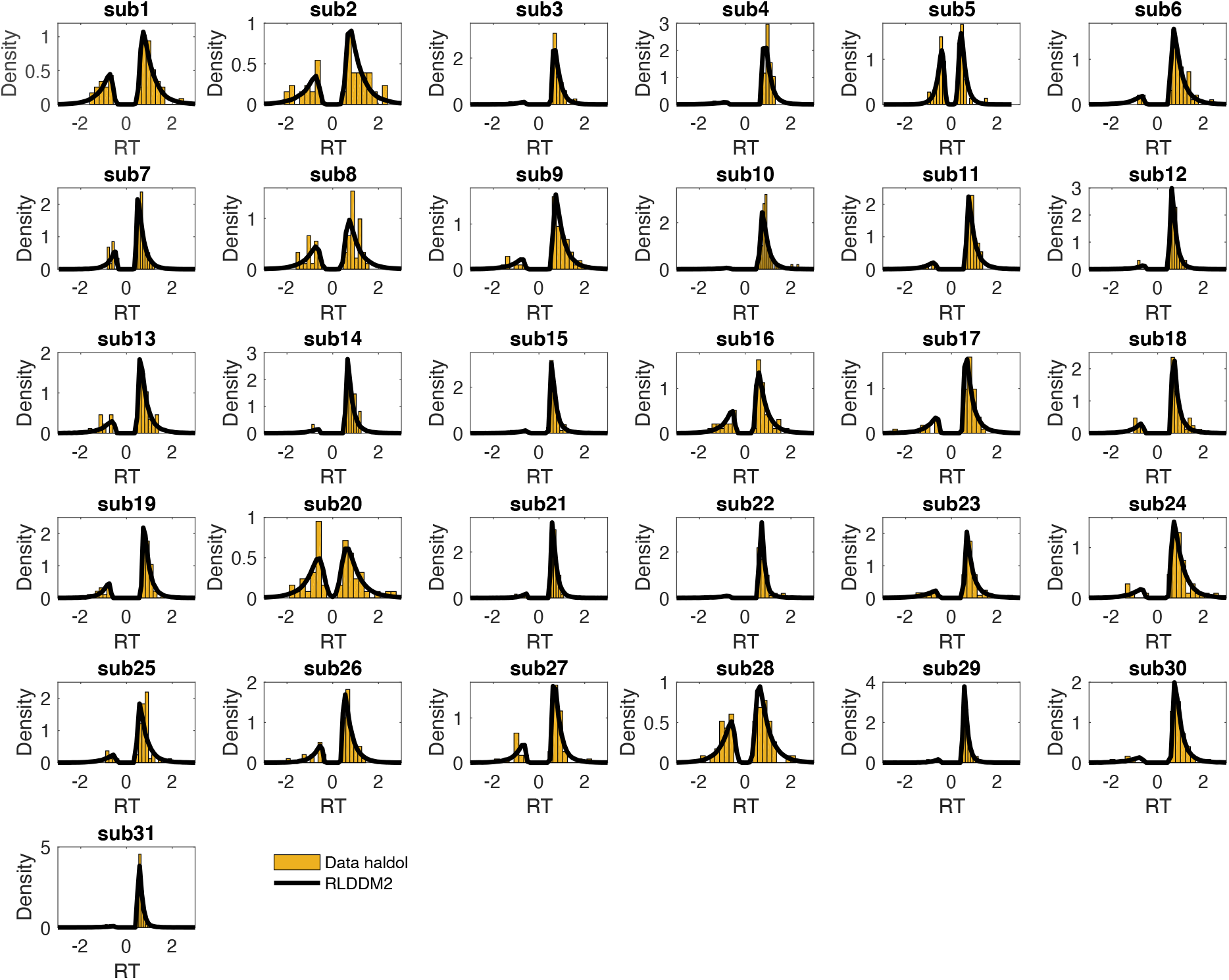
Single-Subject posterior predictive checks for the haloperidol condition. Histograms depict single-subject RT distributions (suboptimal choices are plotted as negative RTs). Solid black lines show smoothed histograms across 1k datasets simulated from the RLDDM2 posterior distribution.

**Supplemental Figure S6.**
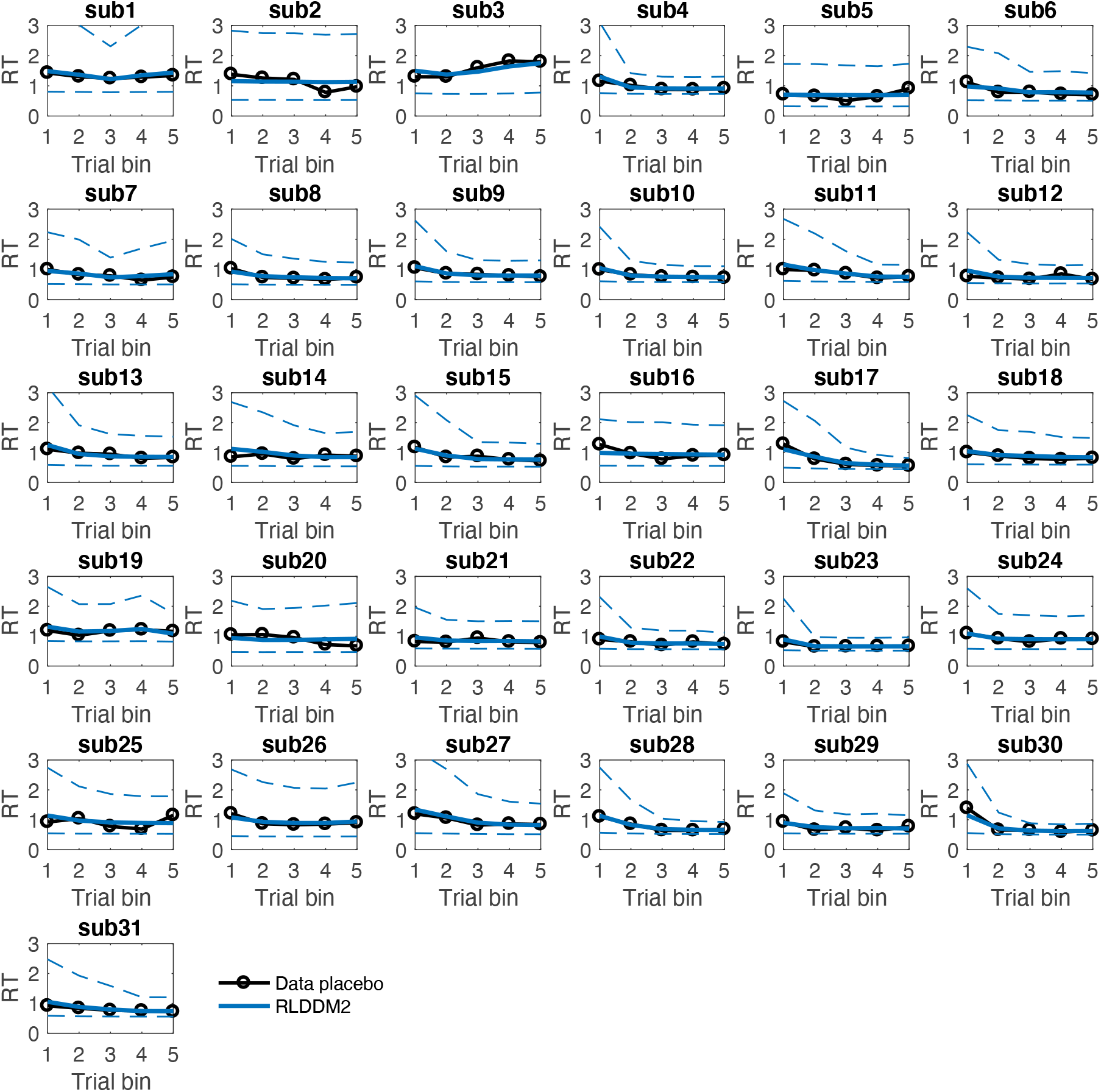
Single-Subject posterior predictive checks of RLDDM2 for RT changes over the course of learning in the placebo condition. Black lines show mean observed RTs per trial bin. Solid colored lines show mean simulated RTs across 1k posterior samples. Dashed lines show +/− 95% percentiles of the simulations.

**Supplemental Figure S7.**
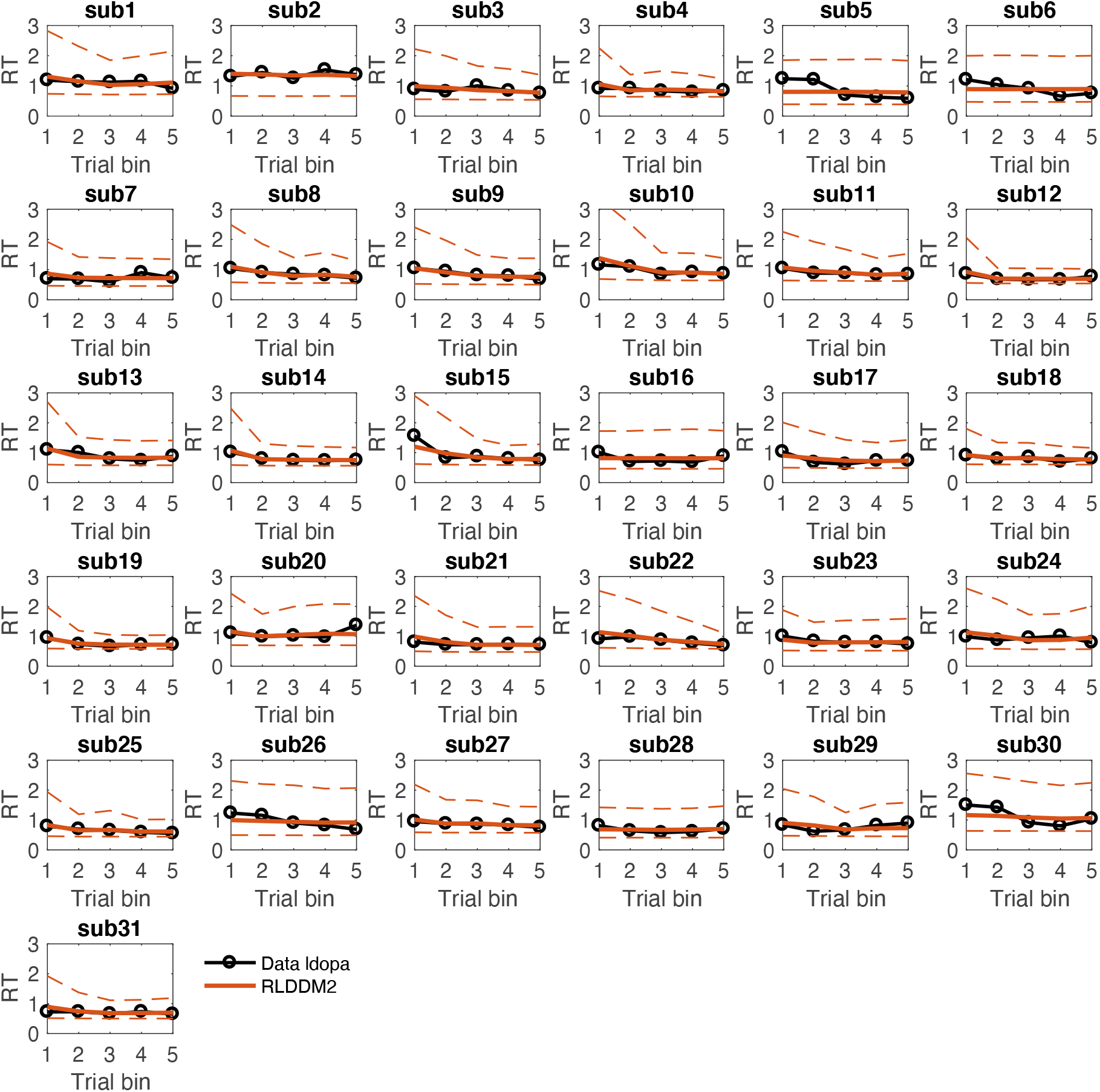
Single-Subject posterior predictive checks of RLDDM2 for RT changes over the course of learning in the L-Dopa condition. Black lines show mean observed RTs per trial bin. Solid colored lines show mean simulated RTs across 1k posterior samples. Dashed lines show +/− 95% percentiles of the simulations.

**Supplemental Figure S8.**
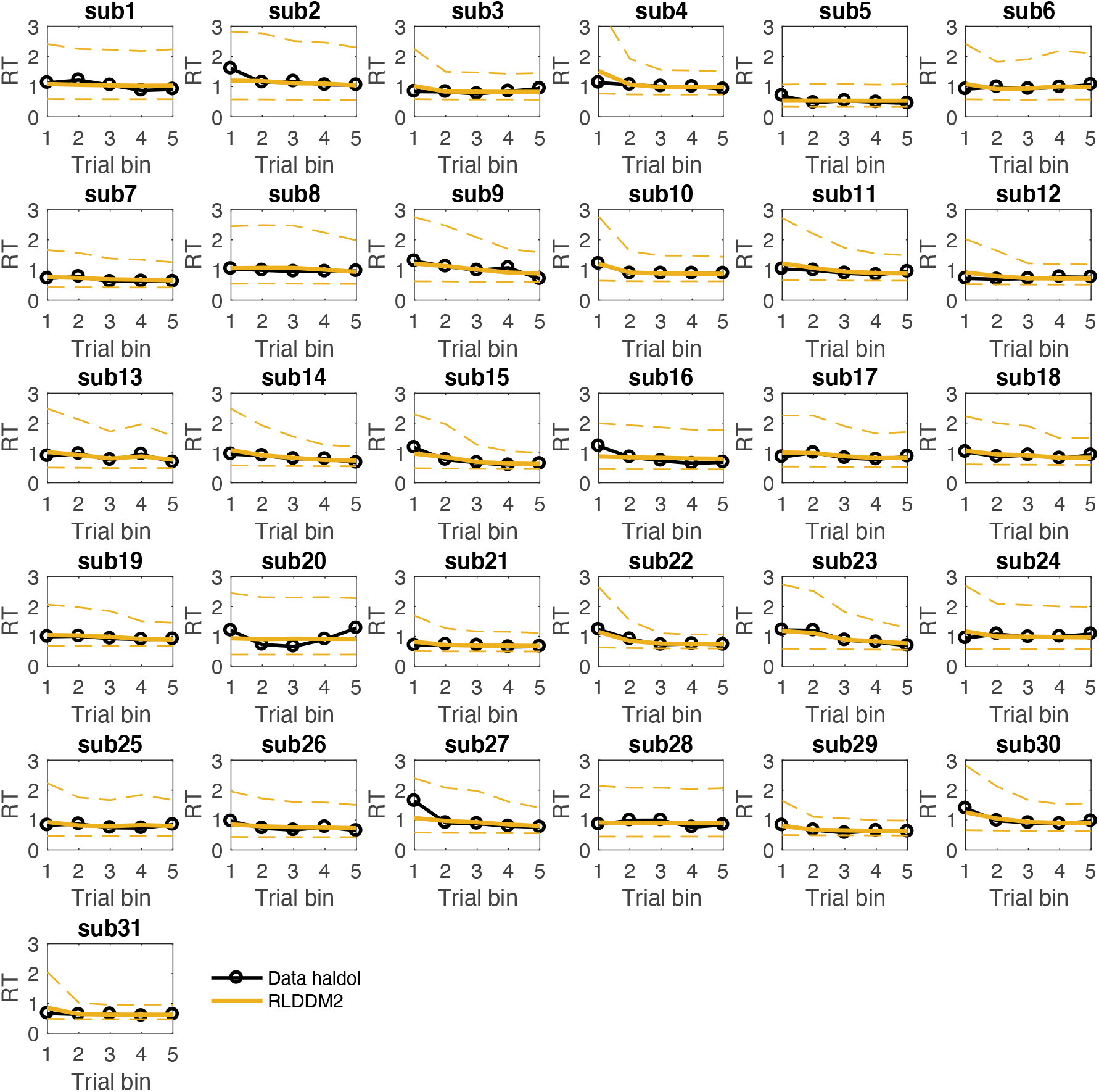
Single-Subject posterior predictive checks of RLDDM2 for RT changes over the course of learning in the haloperidol condition. Black lines show mean observed RTs per trial bin. Solid colored lines show mean simulated RTs across 1k posterior samples. Dashed lines show +/− 95% percentiles of the simulations.

**Supplemental Figure S9.**
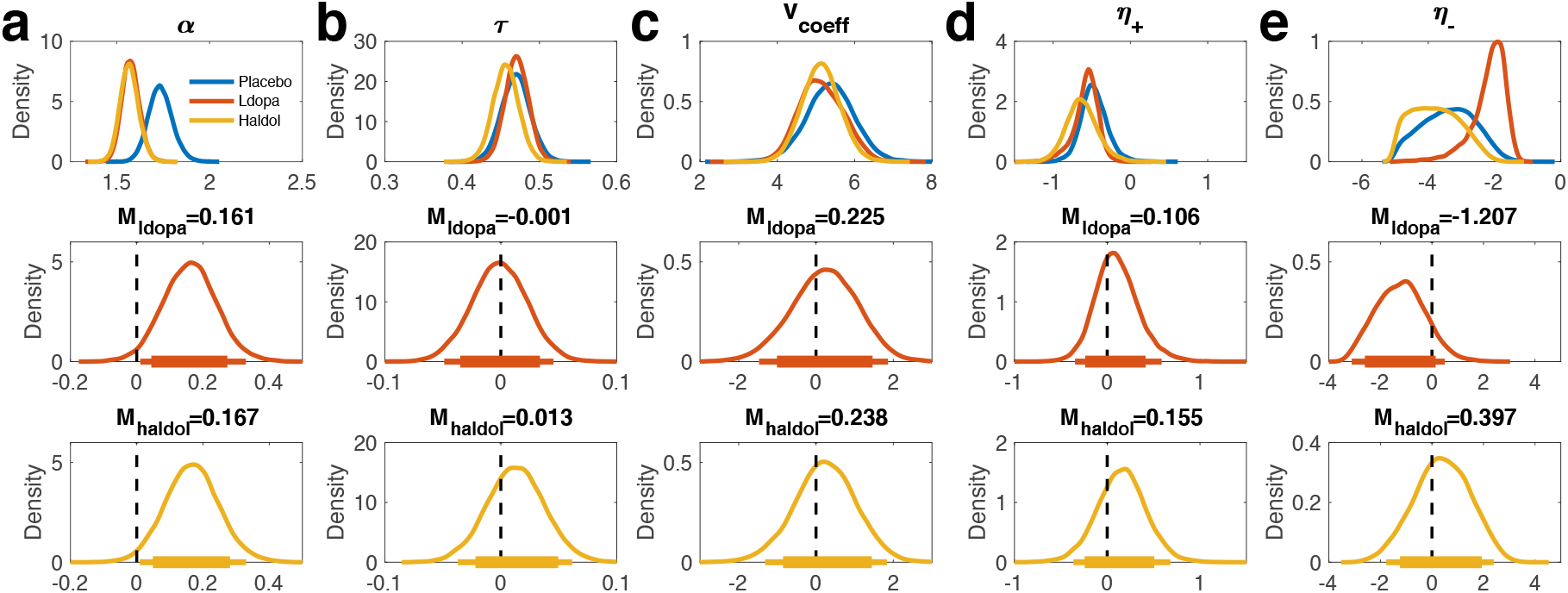
Drug effects on RLDDM parameters when modeling each drug condition separately. a: boundary separation (*α*), b: non-decision time (*τ*), c: value coefficient of the drift rate (*ν*_coeff_), d: positive learning rate (η_+_) in standard normal space, e: negative learning rate (η_−_) in standard normal space. Top row: Posterior distributions per drug condition. Center row: Posterior distribution differences (Placebo – Ldopa, M_ldopa_ refers to the mean of the difference). Bottom row: Posterior distribution differences (Placebo – Haldol, M_haldol_ refers to the mean of the difference). Solid (thin) horizontal lines denote 85% (95%) highest posterior densities.

**Supplemental Table S3.** Mean posterior differences in model parameters (M_diff_) from a modeling scheme in which separate RLDDMs were fit per drug condition. for placebo – ldopa (left) and placebo – haloperidol (right) as well as Bayes factors testing for directional effects (dBF). dBF values > 1 quantify the degree of evidence for a reduction in a parameter compared to placebo, compared to the evidence for an increase. dBF values < 1 reflect the reverse.

| RLDDM<br>parameter | Placebo vs. Ldopa |  | Placebo vs. Haldol |  |
| --- | --- | --- | --- | --- |
| | $M_{\text{diff}}$ | dBF | $M_{\text{diff}}$ | dBF |
| $\alpha$ | .161 | 41.47 | .167 | 48.72 |
| $\tau$ | .001 | .93 | .013 | 2.39 |
| $v_{\text{coeff}}$ | .225 | 1.63 | .238 | 1.65 |
| $\eta_+$ | .106 | 2.16 | .155 | 2.81 |
| $\eta_-$ | -1.207 | .12 | .397 | 1.74 |

